# An antibody-dependent enhancement (ADE) activity eliminated neutralizing antibody with potent prophylactic and therapeutic efficacy against SARS-CoV-2 in rhesus monkeys

**DOI:** 10.1101/2020.07.26.222257

**Authors:** Shuang Wang, Yun Peng, Rongjuan Wang, Shasha Jiao, Min Wang, Weijin Huang, Chao Shan, Wen Jiang, Zepeng Li, Chunying Gu, Ben Chen, Xue Hu, Yanfeng Yao, Juan Min, Huajun Zhang, Ying Chen, Ge Gao, Peipei Tang, Gang Li, An Wang, Lan Wang, Shuo Chen, Xun Gui, Jinchao Zhang, Zhiming Yuan, Datao Liu

## Abstract

Efficacious interventions are urgently needed for the treatment of COVID-19. Here, we report a monoclonal antibody (mAb), MW05, showing high SARS-CoV-2 neutralizing activity by disrupting the interaction of receptor binding domain (RBD) with angiotensin-converting enzyme 2 (ACE2) receptor. Crosslinking of Fc with FcγRIIB mediates antibody-dependent enhancement (ADE) activity by MW05. This activity was eliminated by introducing the LALA mutation to the Fc region (MW05/LALA). Most importantly, potent prophylactic and therapeutic effects against SARS-CoV-2 were observed in rhesus monkeys. A single dose of MW05/LALA completely blocked the infection of SARS-CoV-2 in a study of its prophylactic effect and totally cleared SARS-CoV-2 in three days in a treatment setting. These results pave the way for the development of MW05/LALA as an effective strategy for combating COVID-19.

---

COVID-19, caused by SARS-CoV-2, is currently spreading globally, threatening human health and economic development ^1,2^. As of July 27, 2020, COVID-19 has resulted in more than 16 million infections and 647,784 deaths. Although, multiple clinical trials are ongoing to evaluate repurposing anti-viral and anti-inflammatory agents, no specific treatment against SARS-CoV-2 has been approved since the worldwide outbreak began six months ago ^3^. Treatments using plasma from convalescent COVID-19 patients have shown clear clinical improvement of both mild and severe cases of COVID-19, indicating that passive administration of neutralizing mAbs could have a major impact on controlling the SARS-CoV-2 pandemic by providing immediate protection ^4,5^. During the SARS and Middle East respiratory syndrome coronavirus (MERS-CoV) outbreaks, a number of neutralizing mAbs were developed and proved their potential therapeutic uses for the treatment of coronavirus infections ^6,7^. Neutralizing antibodies for Ebola virus, mAb114 and REGN-EB3, are other encouraging examples that using antibody-based therapy can be effective during an infectious disease outbreak ^8–10^.

The spike (S) protein on the surface of SARS-CoV-2 is the major molecular determinant for viral attachment, membrane fusion and entry into host cells. Therefore, this protein is the main target for development of neutralizing antibodies and vaccines. Previous studies revealed that a large number of antibodies targeting the receptor binding domain (RBD) of either SARS-CoV or MERS-CoV showed potent neutralizing activities by disrupting the interaction of spike protein with receptors on host cells ^11–13^. Screening of RBD targeting antibodies is the most straightforward way to generate SARS-CoV-2 neutralizing antibodies.

To obtain fully human SARS-CoV-2 neutralizing mAbs, we first generated SARS-CoV-2 RBD recombinant protein. We used this protein as bait to isolate specific memory B cells from peripheral blood mononuclear cells (PBMCs) of a COVID-19 convalescent patient. We then used a single B cell cloning strategy to amplify the variable regions of IgG antibodies from individual B cells and insert them into human IgG1 vectors for recombinant antibody expression ^14^. A large panel of SARS-CoV-2 RBD-binding mAbs were generated and characterized. Two mAbs, MW05 and MW07, showed high RBD binding abilities and strong RBD/ACE2 disrupting activities in ELISA. IC_50_ was determined to be 0.054 μg/mL for MW05 and 0.037 μg/mL for MW07. (Fig. 1 A to C). FACS analysis showed that both mAbs could specifically bind to SARS-CoV-2 S protein expressed on HEK293 cells (Fig. 1D). The dissociation constants (K_d_) of MW05 and MW07 binding to SARS-CoV-2 S1 recombinant protein was measured by a surface plasmon resonance (SPR) assay. K_d_ was 0.403 nM for MW05 and 0.462 nM for MW07 (Fig.1E). No cross reactivity with SARS-CoV or MERS-CoV S1 recombinant proteins was detected for either mAb, as assessed using ELISA (Fig. 1 F). SARS-CoV-2 raced around the world after its initial outbreak. Over the past few months it has been mutating. We next expressed RBD recombinant proteins from eight SARS-CoV-2 strains with reported high-frequency mutations. Binding assays showed that both MW05 and MW07 exhibited the same binding abilities to all RBD recombinant proteins. This result suggests that MW05 and MW07 may neutralize all eight of these strains (Fig. 1G; Extended Data Fig. 1).

**Fig. 1.**
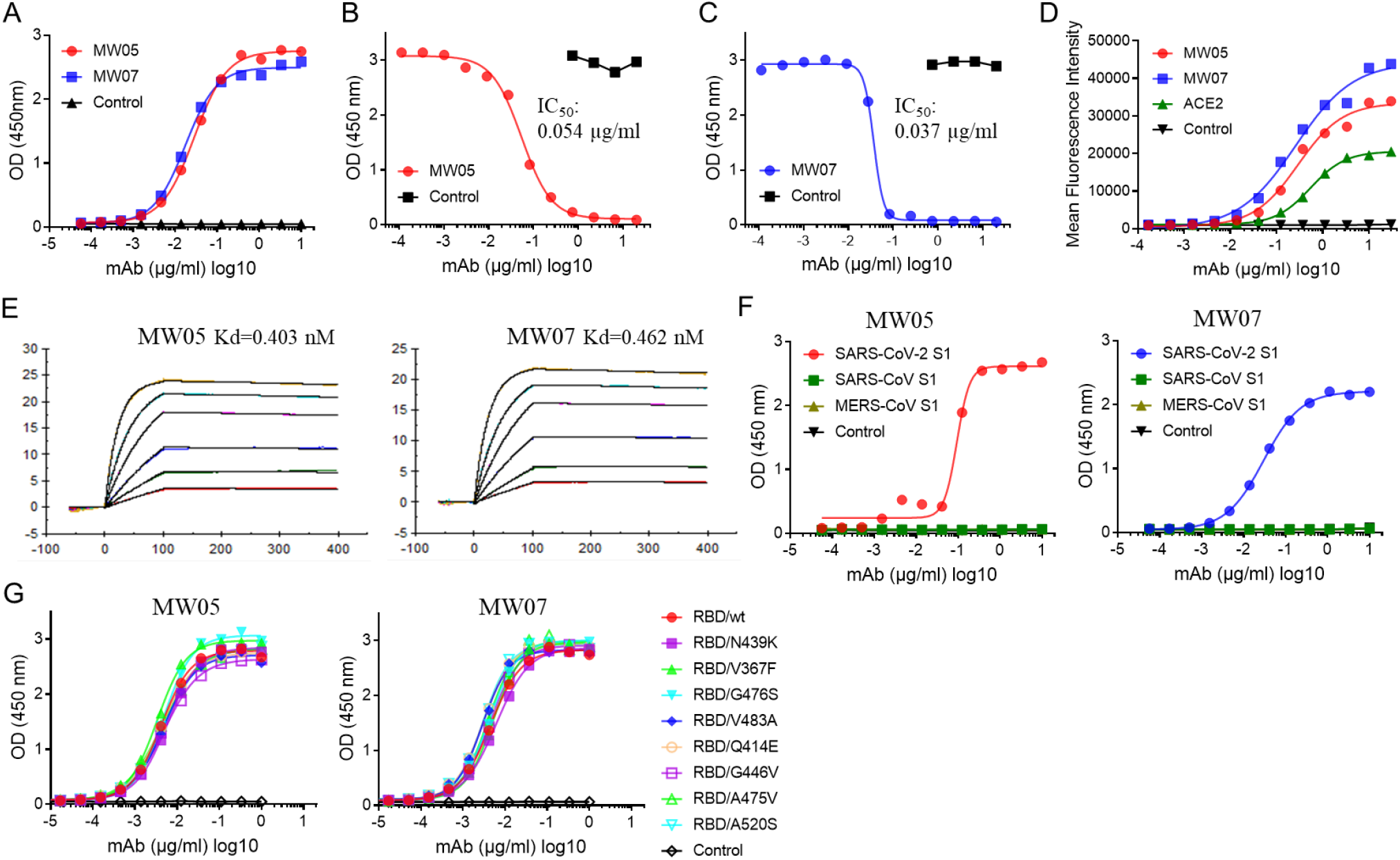
MW05 and MW07 disrupting the interaction of SARS-CoV-2 RBD with hACE2 receptor. (**A**) The binding abilities of MW05 and MW07 to SARS-CoV-2 RBD recombinant protein were assessed by ELISA. (**B** and **C**) The ability of MW05 and MW07 to block SARS-CoV-2 RBD interaction with ACE2 was evaluated by competition ELISA. (**D**) The binding of MW05 and MW07 to SARS-CoV-2 S protein expressed on HEK293 cells was measured by FACS. (**E**) The dissociation constants (K_d_) of MW05 and MW07 to SARS-CoV-2 S1 recombinant protein were measured using a BIAcore S200 system. (**F**) The cross-reactivities of MW05 and MW07 to SARS-CoV-CoV-2, SARS-CoV and MERS-CoV recombinant S1 subunit of spike proteins (S1) were tested by ELISA. (**G**) The binding of MW05 and MW07 to RBD recombinant proteins of SARS-CoV-2 mutated strains.

To investigate the neutralizing activities of MW05 and MW07, we used *in vitro* assays to assess neutralization of first pseudovirus bearing the S protein of SARS-CoV-2 and then authentic virus. Both MW05 and MW07 inhibited pseudovirus infection of Huh7 cells effectively. NT_50_ was measured as 0.030 μg/ml for MW05 and 0.063 μg/ml for MW07 (Fig. 2, A and B). We further evaluated the neutralizing activities of these two mAbs with authentic SARS-CoV-2 infection of Vero E6 cells. As expected, MW05 and MW07 blocked authentic SARS-CoV-2 entry into Vero E6 cells, with 100% neutralization titer (NT_100_) around 1 μg/ml for MW05 and 5 μg/ml for MW07 (Fig. 2, C and D). In summary, MW05 and MW07 exhibited substantial neutralization of both SARS-Cov-2 pseudovirus and authentic virus.

**Fig. 2.**
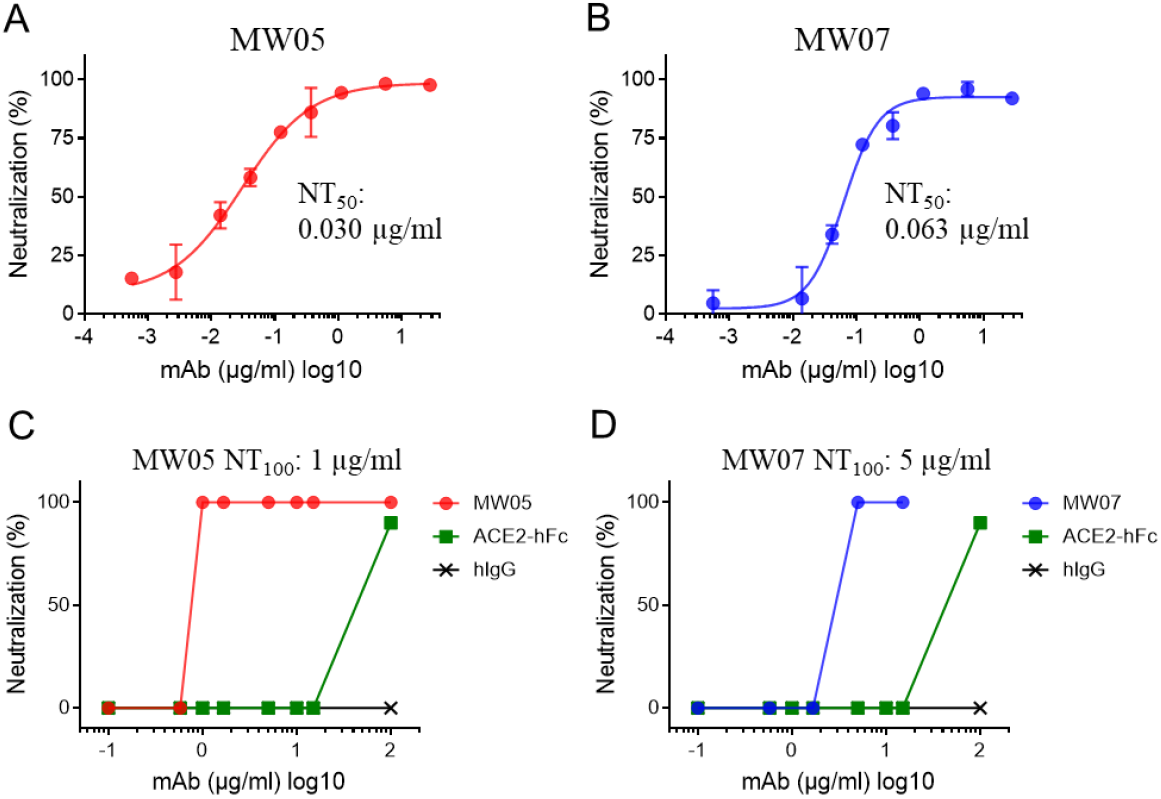
Neutralizing activities of MW05 and MW07. (**A** and **B**) SARS-CoV-2 pseudovirus neutralizing activities of MW05 and MW07 were evaluated on Huh7 cells. 50% neutralization titer (NT_50_) was calculated by fitting the luciferase activities from serially diluted antibodies to a sigmoidal dose-response curve. (**C** and **D**) SARS-CoV-2 authentic virus neutralizing activities of MW05 and MW07 were evaluated using Vero E6 cells. 100% neutralization titer (NT_100_) was labeled accordingly.

ADE has been observed for coronaviruses and several publications have shown that sera induced by SARS-CoV S protein enhanced viral entry into immune cells and inflammation ^15,16^. To evaluate ADE activities of MW05 and MW07, we assessed the infection of SARS-CoV-2 pseudovirus and mAbs complex in THP-1, K562 and Raji cells. These cells are resistant to SARS-CoV-2 pseudovirus infection, as they do not express ACE2 receptor (Extended Data Fig. 2). Cells were incubated with the mixture of pseudovirus with serially diluted MW05. Enhanced SARS-CoV-2 pseudovirus infection of Raji cells, but not of THP-1 o K562 cells was observed (Fig. 3A). Interestingly, No ADE activity was detected for MW07 on all three cell lines (Fig. 3B). Next, we determined the FcγR expression profile of the three cell lines. FACS data revealed that Raji cells, which showed ADE activity for MW05, only express a relatively high level of FcγRIIB; THP-1 cells express high levels of FcγRIA and FcγRIIA; and K562 cells only express high level of FcγRIIA (Fig. 3C). These results indicate that FcγRIIB is the major FcγR contributing to the enhancement of SARS-CoV-2 infection mediated by MW05.

**Fig. 3.**
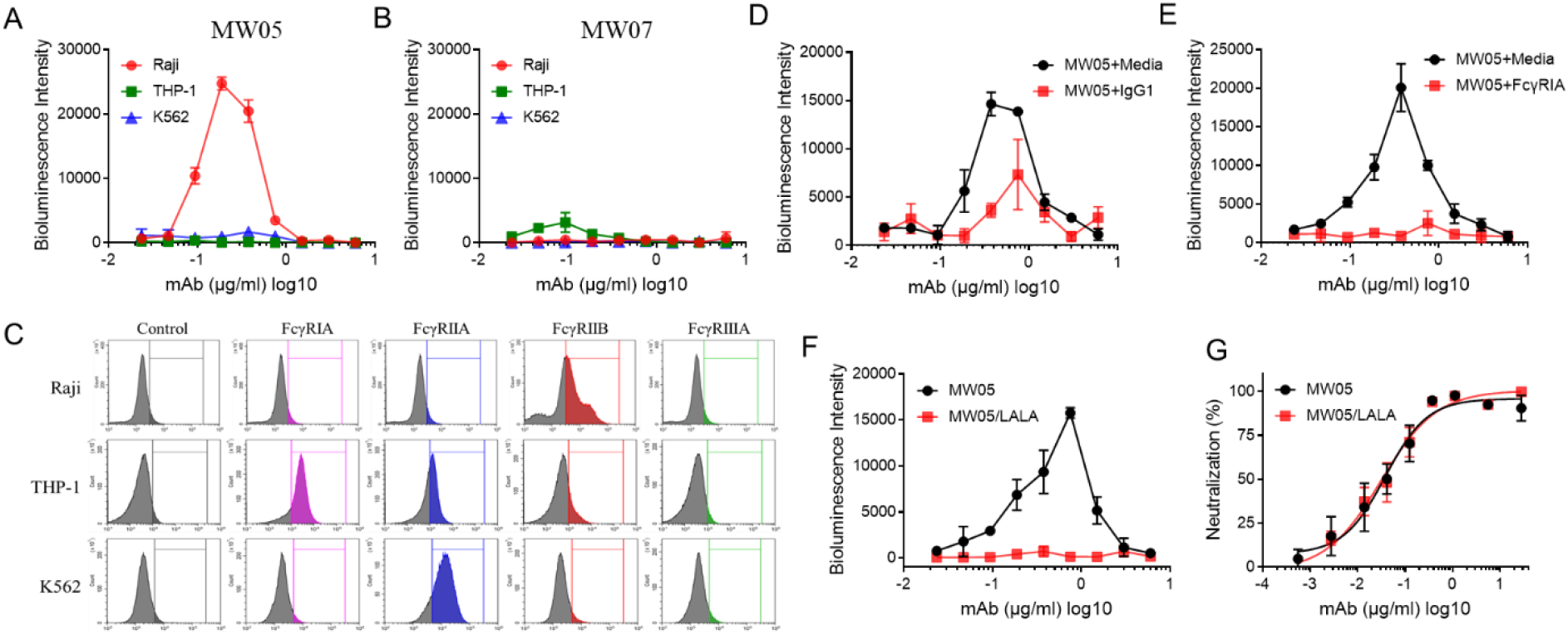
Crosslinking of Fc and FcγR contributing to ADE activities of MW05. (**A** and **B**) ADE activities of MW05 and MW07 were assessed using SARS-CoV-2 pseudovirus. Pseudoviruses pre-incubated with serially diluted mAb mixture were added to Raji, THP-1 and K562 cells to evaluate their ability to enhance infection. RPMI 1640 media containing 10% FBS was used as negative control. (**C**) ADE activities of MW05 on Raji cells pre-treated with media or 400 μg/mL irrelevant hIgG1 were assessed using SARS-CoV-2 pseudovirus. (**D**) ADE activities of MW05 pre-incubated with media or 80 μg/mL FcγRIA were assessed on Raji cells using SARS-CoV-2 pseudovirus. (**E**) The ADE activities of MW05 and MW05/LALA on Raji cells were compared using SARS-CoV-2 pseudovirus. (**F**) The pseudovirus neutralizing activities of MW05 and MW05/LALA on Huh7 cells were measured.

To further assess the ADE activities of MW05, we pre-incubated Raji cells along with irrelevant hIgG1 or MW05 along with FcγRIA recombinant protein to disrupt the interaction of MW05 Fc with FcγRIIB on Raji cells. Both pre-incubation strategies effectively inhibited the ADE activities of MW05 (Fig.3 D and E). FcγRIA has high affinity for the Fc of human IgG1. Accordingly, pre-incubation of MW05 with FcγRIA recombinant protein showed higher ADE inhibition than did pre-incubation of irrelevant hIgG1with Raji cells by disruption of MW05 Fc with with FcγRIIB on Raji cells (Fig.3 D and E). To eliminate the risk of ADE and Fc-mediated acute lung injury *in vivo*, we introduced the LALA mutation to the Fc region of MW05 (MW05/LALA) to decrease the engagement of MW05 with FcγRs. This mutation completely eliminated ADE activity of MW05 without decreasing its neutralizing activity (Fig. 3 F and G).

We evaluated the prophylactic and therapeutic effects of MW05/LALA in a rhesus monkey SARS-CoV-2 infection model. In the prophylactic (pre-challenge) group, three animals were injected intravenously with a single dose of MW05/LALA (20 mg/kg) one day before receiving a 1×10^5^ 50% tissue culture infectious dose (TCID_50_) SARS-CoV-2 challenge via intratracheal incubation (Fig. 4A). MW05/LALA antibody effectively protected animals from SARS-CoV-2 infection; almost no virus was detected in the oropharyngeal swabs of the prophylactic group (Fig. 4B).

**Fig. 4.**
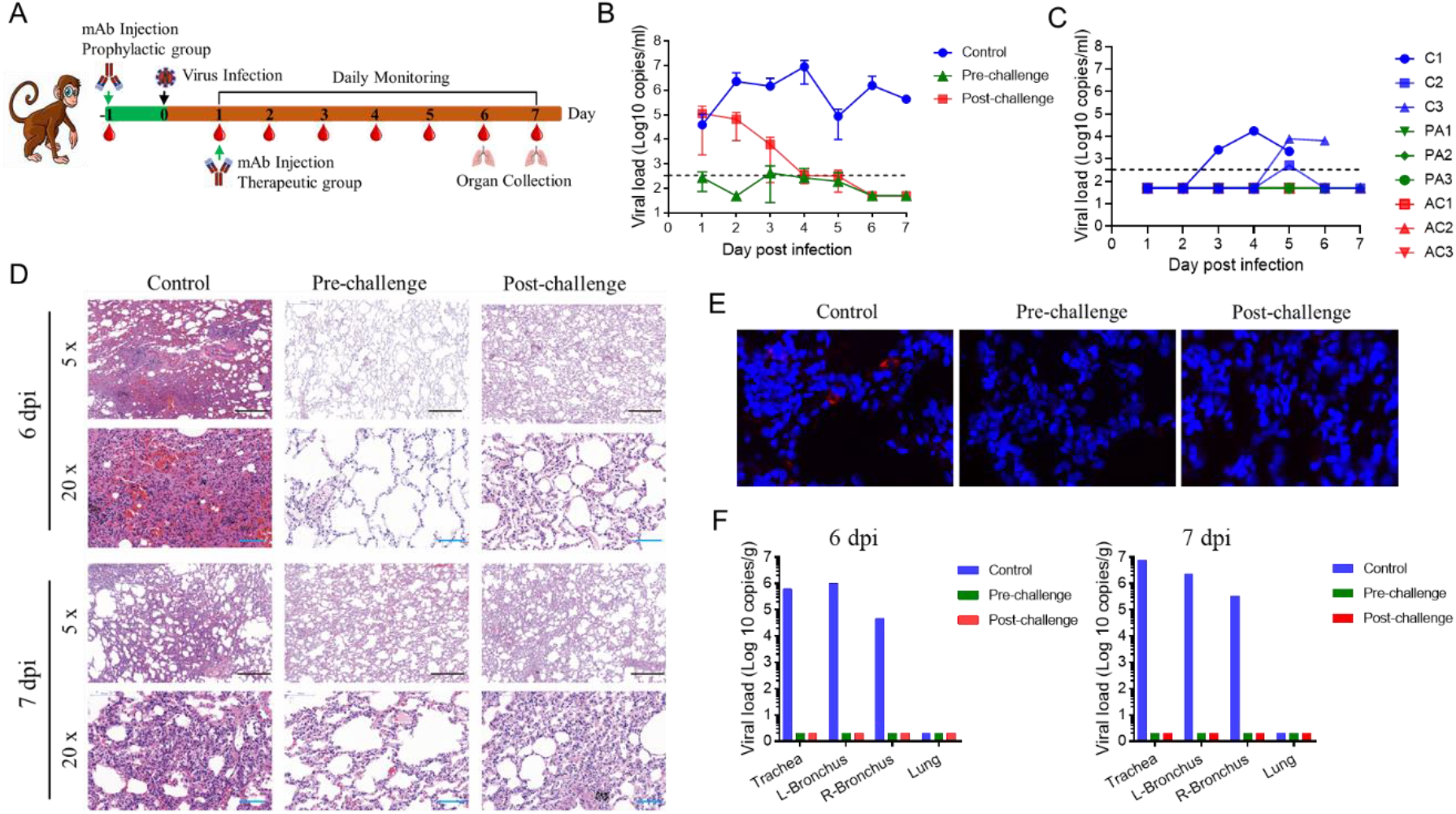
Prophylactic and therapeutic effects of MW05/LALA. (**A**) A schematic of the experimental *in vivo* set up. Nine rhesus monkeys were divided into pre-challenge (prophylactic), post-challenge (therapeutic) and control groups with 3 animals in each group. Before virus challenge, the monkeys in the pre-challenge group were injected intravenously with a single dose of 20 mg/kg MW05/LALA. One day later, all monkeys were challenged with 1×10 ^5^ TCID_50_ SARS-CoV-2 via intratracheal intubation. A single dose of 40 mg/kg MW05/LALA was administered to each animal in the post-challenge group on day 1 post challenge. Monkeys in the control group were given 20 mg/kg irrelevant hIgG1 one day before virus challenge. (**B**) Viral titer of oropharyngeal swabs at the indicated time points were evaluated using qRT-PCR. Data are average values from three monkeys (n=3) for the first 5 days, from two monkeys (n=2) for 6 dpi, and from one monkey (n=1) for 7 dpi. The line for limit of detection is labeled. (**C**) Viral titer of rectal swabs at the indicated time point were evaluated by qRT-PCR. “C” indicates the control group, “PA” indicates the pre-challenge group and “AC” indicates the post-challenge group. (**D**) Histopathology and immunohistochemical examination of lung tissues from pre-challenge, post-challenge and control monkeys. (**E**) Immunohistochemical analysis of SARS-CoV-2 protein expression in lung tissues from pre-challenge, post-challenge and control monkeys. (**F**) Viral load analysis of trachea, bronchus and lung tissues of experimental animals. L-Bronchus means left bronchus; R-Bronchus means right bronchus.

In the treatment (post-challenge) group, three animals were first challenged with 1×10^5^ TCID_50_ SARS-CoV-2. Then, at day 1 post infection (dpi), a single dose of MW05/LALA (40 mg/kg) was administered intravenously to these animals (Fig. 4A). Animals in the control group (n=3) were given a single dose of irrelevant hIgG1 (20 mg/kg) on 1 dpi. In the control group, the viral loads in oropharyngeal swabs increased to a peak of about 10^7.0^ RNA copies/mL on 4 dpi, then decreased to the limit of detection on 7 dpi (Fig. 4B). Virus was only detected in the rectal swabs of two animals in the control group (Fig. 4C). Notably, virus titers decreased in the MW05/LALA treatment group immediately after administration. No virus was detected in the MW05/LALA treatment group even on 4 dpi, the time point at which viral titers in the control group reached their peak. A single dose of MW05/LALA exhibited SARS-CoV-2 therapeutic efficacy in a rhesus monkey model, clearing virus in three days after antibody administration (Fig. 4B). No significant weight loss or body temperature change was observed in any of the animals during the study (Extended Data Fig. 3 and 4). No virus was detected in nasal swabs or blood samples (Extended Data Fig. 5). Additionally, no significant abnormal hematology changes were observed (Extended Data Fig. 6).

Rhesus monkeys challenged with SARS-CoV-2 were evaluated for tissue damage. One monkey from each group was euthanized for necropsy on 6 and 7 dpi. Interstitial pneumonia symptoms were observed in the control group, including thickened alveolar septa, intensive infiltration of monocytes and lymphocytes, and proliferation of fibroblasts (Fig. 4D). We also observed cellulose exudation in some alveolar cavities, with the formation of hyaline membrane and pulmonary hemorrhaging (Fig. 4D). Monkeys in treatment group displayed limited pathological lung changes, with overall alveolar structure intact and much lower levels of fibroblasts proliferation and leukocyte infiltration than were observed in the control monkeys (Fig. 4D). No lesions were observed in the lungs of the animal euthanized on 6 dpi and very mild pulmonary hemorrhaging of the animal euthanized on 7 dpi in the prophylactic group (Fig. 4D). In summary, MW05/LALA effectively inhibited lung tissue damage in both prophylactic and therapeutic ways in a rhesus monkey SARS-CoV-2 infection model.

Immunohistochemical analysis of virus in lung tissues showed that SARS-CoV-2 protein only been detected in the lung tissue of the control group on 5 dpi but not 6 dpi and 7 dpi. In comparison, viral proteins were undetectable in the lung tissues of animals in the prophylactic and therapeutic groups (Fig. 4E). In order to further understand the distribution of SARS-CoV-2 in upper respiratory tract, trachea and bronchus tissue samples were collected on 6 dpi and 7 dpi. Viral titers were then determined by qRT-PCR. On 6 dpi and 7 dpi, high levels of SARS-CoV-2 RNA copies were detected in trachea and bronchi tissues of control animals, while no viral nucleic acid was detected from tissue samples of both prophylactic and therapeutic groups (Fig. 4F).

The global COVID-19 pandemic is running rampant over the world. There are great unmet medical needs for COVID-19 therapy as no SARS-CoV-2 specific drugs or vaccines have yet been approved. Neutralizing mAbs are promising agents to combat emerging infectious diseases. Our results showed the prophylactic and therapeutic efficacy of MW05/LALA on SARS-CoV-2 *in vivo*. This work paves the way for further development of antibody-based therapies for prophylactic or therapeutic treatment of COVID-19.

## Methods

### Ethics statement

All neutralizing assays using SARS-CoV-2 authentic virus were performed in biosafety level 3 (BSL-3) facility. Monkey studies were carried out in an animal biosafety level 4 (ABSL-4) facility with protocols approved by the Laboratory Animal Welfare and Ethics Committee of the Chinese Academy of Sciences. The blood was taken from a convalescent COVID-19 patient after got his signature for the informed consent form.

### Cells and viruses

HEK293 (ATCC, CRL-3216) cells, Huh7 (Institute of Basic Medical Sciences CAMS, 3111C0001CCC000679) cells and Vero E6 (ATCC, CRL-1586) cells were cultured at 37 °C in Dulbecco’s Modified Eagle medium (DMEM) supplemented with 10% fetal bovine serum (FBS). Raji (ATCC, CCL-86) cells, THP-1 (ATCC, TIB-202) cells and K562 (ATCC, CCL-243) cells were cultured at 37 °C in RPMI 1640 Medium with 10% FBS. SARS-CoV-2 was isolated by the Center for Disease Control and Prevention of Zhejiang province. Vero E6 cells were applied to the reproduction of SARS-CoV-2 stocks.

### Recombinant protein generation

The SARS-CoV-2 RBD (319-533aa, accession number: QHD43416.1), SARS-CoV-2 S1 (1-685aa, accession number: QHD43416.1) and SARS-CoV-2 RBD mutants recombinant proteins tagged with C-terminal 6 × His were cloned into the pKN293E expression vector. HEK293 cells were transiently transfected with plasmids using 293fectinTM Transfection Reagent (Cat: 12347019, Life Technologies) when the cell density reached 1 × 10^6^ cells/mL. Four days after transfection, the conditioned media was collected by centrifugation followed by purification using HisTrapTM HP (Cat: 17-5248-01, GE Healthcare). The purified protein was buffer exchanged into PBS using a Vivacon 500 concentrator (Cat: VS0122, Sartorius Stedim). For the generation of human ACE2-hFc and SARS-CoV-2 RBD-mFc recombinant proteins, RBD or ACE2 sequence (1-615aa, accession number: NP_068576.1) was cloned into mouse IgG1 or human IgG1 Fc backbone in pKN293E expression vectors and transiently transfected into HEK293 cells followed by media collection and purification using MabSelect SuRe antibody purification resin (Cat: 29-0491-04, GE Healthcare). SEC-HPLC and SDS-PAGE were used to check the size and purity of these recombinant proteins.

SARS-CoV-2 RBD mutant information:

| Mutants | Virus Strain Name | Accession ID | Data Source |
| --- | --- | --- | --- |
| RBD/NE39K | hCoV-19/Scotland/EDB162/2020 | EPI_ISL_425924 | GISAID |
| V367F | HCoV-19/England/20134027504/2020 | EPI_ISL_423136 | GISAID |
| G476S | hCoV-19/USA/WA-S28/2020 | EPI_ISL_417081 | GISAID |
| V483A | hCoV-19/USA/WA-S529/2020 | EPI_ISL_434289 | GISAID |
| Q414E | hCoV-19/USA/AZ-TGEN-TG268099/2020 | EPI_ISL_426500 | GISAID |
| G446V | hCoV-19/Australia/VIC329/2020 | EPI_ISL_426639 | GISAID |
| A475V | hCoV-19/USA/AZ-TGEN-TG268282/2020 | EPI_ISL_426504 | GISAID |
| A520S | hCoV-19/USA/WA_0432/2020 | EPI_ISL_426441 | GISAID |

### Antibody discovery and expression

mAbs were generated from SARS-CoV-2 RBD specific memory B cells using single B cell isolation and cloning strategy ^17^. For preparation of MW05 and MW07 recombinant antibodies, heavy chain and light chain plasmids were transiently co-transfected into HEK293 cells or stably expressed in CHO cells followed by purification with Protein A resin. Antibodies MW05/LALA and MW07/LALA were generated by introducing the LALA mutation (L234A and L235A) in the Fc region of IgG1 to abolish binding with FcγRs and prepared using the same protocol used for generation of wild-type mAbs.

### ELISA

To access the binding of mAbs to recombinant proteins (SARS-CoV-2 RBD, SARS-CoV-2 RBD mutants, SARS-CoV-2 S1, SARS-CoV S1 (Cat: 40150-V08B1, Sino Biological), MERS-CoV S1 (Cat: 40069-V08B1, Sino Biological)), the proteins were first coated on 96 well ELISA plates at 1 μg/ml in 100 μL at 4°C overnight. After blocking with 5% BSA in PBS, serially diluted mAbs were added to the plates and incubated for 60 min at 37°C. Plates were washed and secondary Ab Goat Anti-Human IgG Fc-HRP (Cat: 109-035-098, Jackson ImmunoResearch) was added. TMB was used for color development and absorbance at 450 nm was measured using a microplate reader. For the RBD/ACE2-hFc blocking assay, ACE2-hFc recombinant protein was coated on a 96 well ELISA plate at 0.75 μg/ml in 100 μL at 4°C overnight. Equal volumes (100μL + 100μL) of pre-incubated RBD-mFc/mAb complex (RBD-mFc concentration: 100 ng/ml, mAb concentrations between 40 to 0.00023 μg/ml) were added to the plates and incubated for 60 min at 37°C. Plates were washed and secondary Ab Goat Anti-Mouse IgG Fc-HRP (Cat:115-035-071, Jackson ImmunoResearch) was added. TMB was used for color development and absorbance at 450 nm was measured using a microplate reader.

### Flow cytometry assay

The binding of MW05 and MW07 to S protein expression on cell surface was assessed by FACS. HEK293 cells were transiently transfected by SARS-CoV-2 Spike expression plasmid (Cat: VG40589-UT, Sino Biological) for 24 to 48 hours. Cells were then collected and blocked with 5% BSA for 30 min at RT. 3-fold serially diluted MW05, MW07, ACE2-hFc and isotype control antibody were added into cells (2 × 10^5^ cells/sample in 100 μL) and incubated for 60 min on ice. After washing twice with 1 × PBS, cells were stained with 1/200 diluted Goat Anti human IgG Fc-FITC antibody (Cat: F9512, Sigma) for 45 min and analyzed using flow cytometry (CytoFLEX, Beckman Coulter). The FcγR expression profiles of Raji, THP-1 and K562 were determined by FACS. Cells were collected and washed twice with 1×PBS, then blocked with Fc receptor blocking solution buffer (Cat: MX1505, Maokang Biological) for 30 min at RT. Then 10 μL anti-FcγRI antibody-FITC (Cat: 10256-R401-F, Sino Biological), anti-FcγRIIa antibody-FITC (Cat: 10374-MM02-F, Sino Biological), anti-FcγRIIIa antibody-FITC (Cat: 10389-MM41-F, Sino Biological) and FITC-labeled anti-FcγRIIb antibody (Cat: NBP2-14905, Biotechne; Cat: MX488AS100-1KT, Sigma-Aldrich) were added into cells (1×10^6^ cells/sample in 100 μL) and incubated for 60 min at 2-6°C and analyzed using flow cytometry (CytoFLEX, Beckman Coulter).

### Surface plasmon resonance (SPR)

SPR measurements were performed at room temperature using a BIAcore S200 system with CM4 biosensor chips (GE Healthcare). For all measurements, a buffer consisting of 150 mM NaCl, 10 mM HEPES, 3 mM EDTA, pH 7.4 and 0.005% (v/v) Tween-20 was used as running buffer. All proteins were exchanged into this buffer in advance. The blank channel of the chip served as the negative control. SARS-CoV-2 S1 recombinant protein was captured on the chip at 175 response units. Gradient concentrations of MW05 Fab or MW07 Fab (from 200 nM to 6.25 nM with 2-fold dilution) were then flowed over the chip surface. After each cycle, the sensor was regenerated with Gly-HCl (pH 1.5). The affinity was calculated using a 1:1 (Rmax Local fit) binding fit model with BIAevaluation software.

### Neutralization assay

SARS-CoV-2 pseudovirus was prepared and provided by the Institute for Biological Product Control, National Institutes for Food and Drug Control (NIFDC) ^18^. The TCID_50_ was determined by the transduction of pseudovirus into Huh7 cells. For pseudovirus neutralization assay, 100 μL of mAbs at different concentrations were mixed with 50 μL supernatant containing 500 TCID_50_ pseudovirus. The mixture was incubated for 60 min at 37 °C, supplied with 5% CO_2_. All mAbs were tested in concentrations ranging from 0.55 ng/mL to 28 μg/mL in the context of Huh7 cells. 100 μL of Huh7 cell suspension (2 × 10^5^ cells/mL) was then added to the mixtures of pseudoviruses and mAbs for an additional 24 h incubation at 37 °C. Then, 150 μL of supernatant was removed, and 100 μL luciferase detecting regents (Promega) was added to each well. After 2 mins incubation, each well was mixed 10 times by pipetting, and 150 μL of the mixture was transferred to a new microplate. Luciferase activity was measured using a microplate luminometer (ThermoFisher). The 50% neutralization titer (NT_50_) was calculated using GraphPad Prism 7.0. For SARS-CoV-2 authentic virus neutralization assay, Vero E6 cells were diluted and seeded into a 96-well plate with 1×10^4^ cells/well in 100 μL volume at 37 °C. 16 h later, cells were washed by 1× PBS for 3 times and added diluted antibodies in equal volume with the concentration ranging from 0.1 μg/mL to 100 μg/mL. 100 TCID_50_ SARS-CoV-2 authentic virus was used for each well. Meanwhile, a control group without antibody was set up. A virus back-titration was performed to assess the correct virus titer used in each experiment. Cytopathic effect of each well was monitored every day and photographed at day 3 or day 4 after virus infection. All experiments were conducted following the standard operating procedures of the approved BSL-3 facility.

### Antibody-dependent enhancement (ADE) assay

The ADE assays were performed using Raji, THP-1 and K562 cell lines. 25 μL of 2-fold serially diluted mAbs were mixed with 25 μL supernatant containing 250 TCID_50_ pseudovirus. The mixture was incubated for 60 min at 37 °C, supplied with 5% CO_2_. All mAbs were tested in the concentrations ranging from 6000 to 23.4 ng/mL. 100 μL of THP-1, Raji and K562 cells at the density of 2 × 10^6^ cells/mL were added to the mixtures of pseudoviruses and mAbs for an additional 24 h incubation. Then, same volume of luciferase detecting regents (Promega) was added to each well. After 2 mins incubation, the luciferase activity was measured using a microplate luminometer (Thermo Fisher).

### Animal experiments

All animal experiments were performed according to the procedures approved by the Chinese Academy of Sciences and complied with all relevant ethical regulations regarding animal research. Nine 6 or 7 year-old rhesus monkeys (3 females and 6 males) were divided into 3 groups: a control group (one female and two males), a pre-exposure group (one female and two males) and a post-exposure group (one female and two males). Rhesus monkeys in the control group were injected with 20 mg/kg negative control antibody. For the prophylactic study, monkeys in the pre-exposure group were given a single dose of 20 mg/kg MW05/LALA antibody intravenously one day before being challenged with 1×10^5^ TCID_50_ SARS-CoV-2 via intratracheal routes. For the therapeutic study, monkeys in the post-exposure group were administrated with a single dose of 40 mg/kg MW05/LALA antibody intravenously one day after challenged with 1×10^5^ TCID_50_ SARS-CoV-2 via intratracheal routes. Body weight and body temperature were monitored every day. Oropharyngeal, nasal and rectal swabs were collected for 7 days. Blood samples were collected. White blood cells (WBC), neutrophils (NEUT), lymphocytes (LYMPH) and monocytes (MONO) were assessed for all monkeys. Swabs were placed into 1 mL of DMEM after collection. Viral RNA was extracted by the QIAamp Viral RNA Mini Kit (Qiagen) according to the manufacturer’s instructions. RNA was eluted in 50 μL of elution buffer and used as the template for RT-PCR. The pairs of primers were used targeting S gene: RBD-qF1: 5’-CAATGGTTTAACAGGCACAGG-3’; RBD-qR1: 5’-CTCAAGTGTCTGTGGATCACG-3’. 2 μL of RNA were used to verify the RNA quantity by HiScript^®^ II One Step qRT-PCR SYBR^®^ Green Kit (Vazyme Biotech Co., Ltd) according to the manufacturer’s instructions. The amplification was performed as follows: 50°C for 3 min, 95°C for 30 s followed by 40 cycles consisting of 95°C for 10 s, 60°C for 30 s, and a default melting curve step in an ABI step-one machine (*14*).

### Histopathology and Immunohistochemistry

Animal necropsies were performed according to a standard protocol. Samples for histological examination were stored in 10% neutral-buffered formalin for 7 days, embedded in paraffin, sectioned and stained with hematoxylin and eosin or Masson’s trichrome prior to examination by light microscopy.

### Data availability

Further information and requests for resources and reagents should be directed to and will be fulfilled by the corresponding author Xun Gui.

## Acknowledgements

We thank all colleagues from National Biosafety Laboratory (Wuhan), CAS for their support during the study. We thank Dr. Zhou Yi-Wu for his assistance in the histopathological analysis. We are grateful to Dr. Dayong Tian and Dr. Qi An from Shanghai King-cell Biotechnology Co. Ltd. for helping us performing the SARS-CoV-2 authentic virus neutralizing assays in the P3 laboratory. We thank Ms. Hongyuan Ren and Mr. Pan Gao for their helping in measuring the affinities in SPR assay. We thank Dr. Yu Mao and Ms. Wenlu Liang for the stimulating discussions.

## Funding

This work was supported by National Key R&D Program (2020YFC0848600).

## Author contributions

D.L, J.Z, X.G. and S.W. initiated and coordinated the project. R.W. and S.J. analyzed antigen sequences, expressed and purified recombinant proteins. Z.L., M.W., and P.T. expressed and purified antibodies. R.W. and S.J. performed the binding and blocking assays. B.C. and W.J. performed the pseudovirus neutralization assays. C.G. and W.J. checked the expression profile of FcγRs. W.H. and L.W. designed and supervised the SARS-CoV-2 pseudovirus tests. X.G., B.C., W.J. and C.G. designed and performed the ADE experiments. G.L, A.W. and B.C supervised the protein quality control work. Z.Y., Y.P., C.S., X.H., Y.Y. carried out the monkey studies with the help from H.Z., Y.C. and G.G. J.M. and Y.C. performed histopathology and immunohistochemistry assays. S.C performed the Sars-CoV-2 mutants sequence data analysis and design work. X.G., S.W., S.C. and J.Z. analyzed and prepared the manuscript with input from all authors.

## Competing interests

X.G., S.W., R.W., S.J., W.J. and C.G. are listed as inventors on the licensed patents for MW05 and MW07. S.W., R.W., S.J., M.W., W.J., Z.L., C.G., B.C., P.T., J.Z., X.G. and D.L. are employees of Mabwell (Shanghai) Bioscience Co., Ltd. and may hold shares in Mabwell (Shanghai) Bioscience Co., Ltd. The other authors declare no competing interests.

---

**Extended Data Fig. 1.**
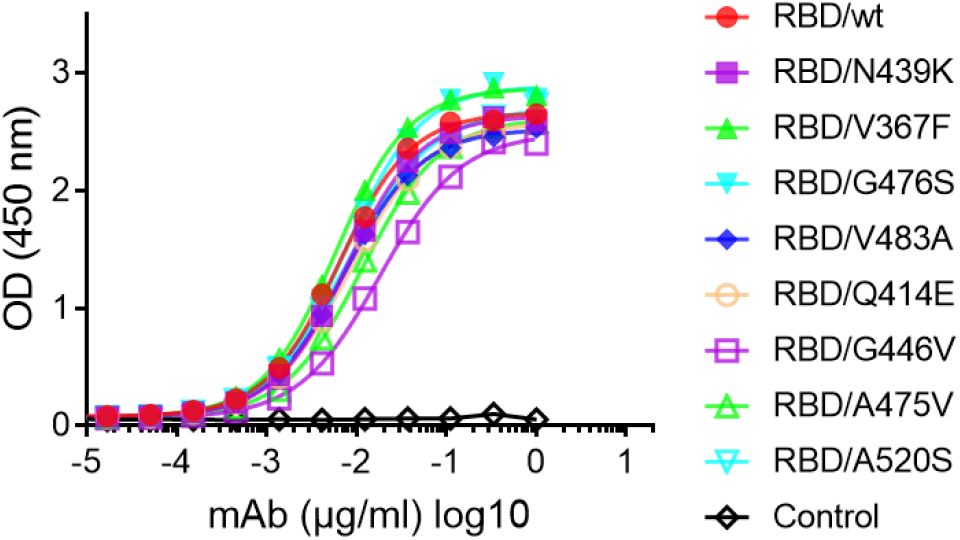
Binding of ACE2 to different SARS-CoV-2 RBD mutants by ELISA. RBD recombinant proteins were coated on 96-well plates. Human ACE2-mFc was then added into plates to check the binding.

**Extended Data Fig. 2.**
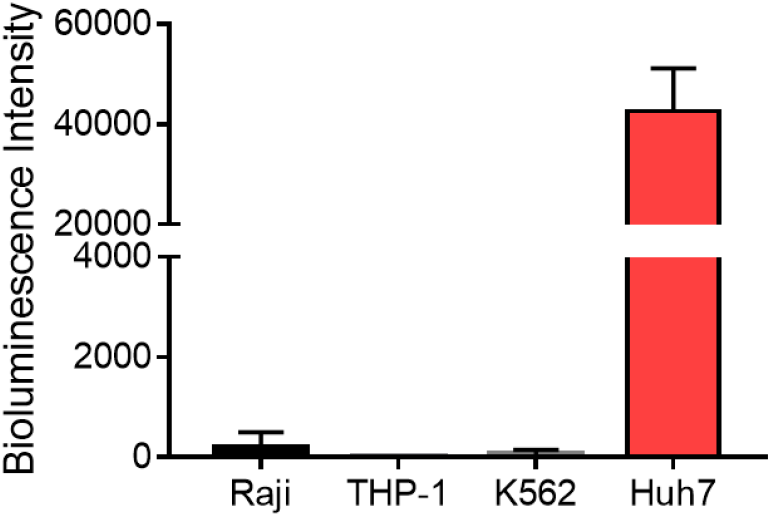
The infection of SARS-CoV-2 pseudovirus in Raji, THP-1, K562 and Huh7 cells.

**Extended Data Fig. 3.**
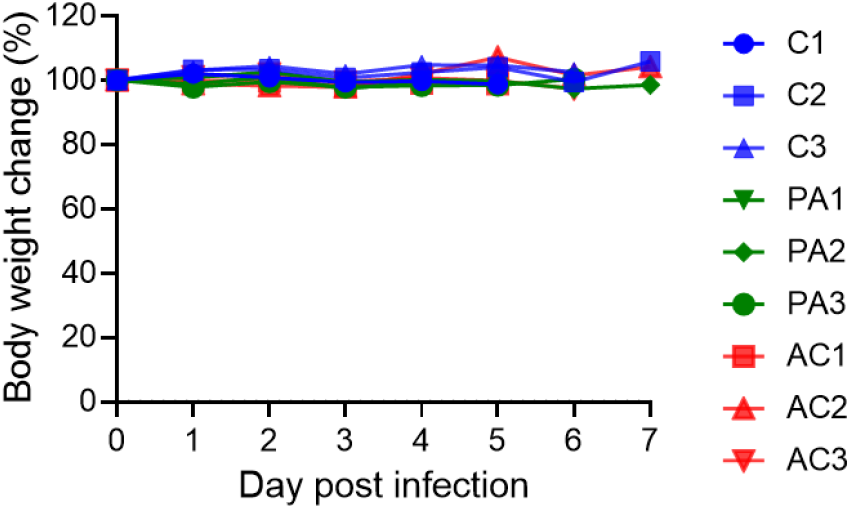
The body weight of each monkey was checked every day. “C” indicates the control group, “PA” indicates the pre-challenge group and “AC” indicates the post-challenge group.

**Extended Data Fig. 4.**
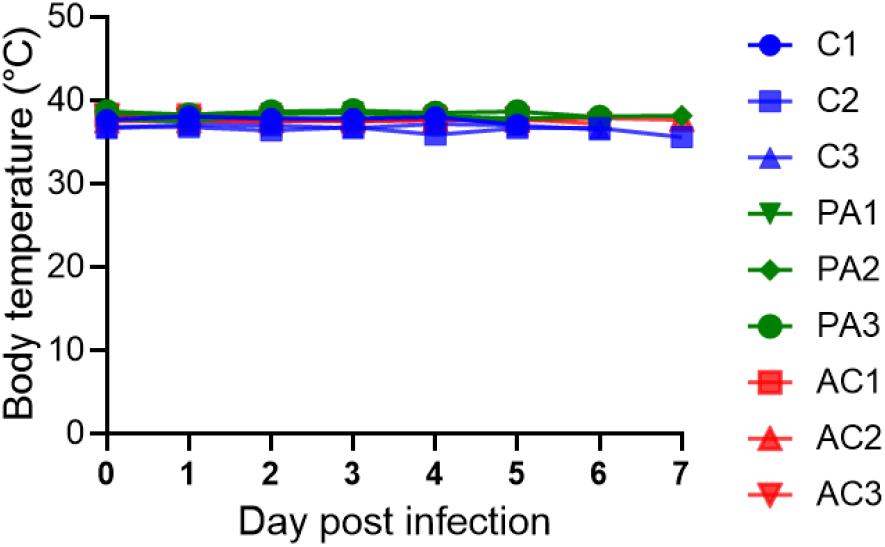
The body temperature of each monkey was checked every day. “C” indicates the control group, “PA” indicates the pre-challenge group and “AC” indicates the post-challenge group.

**Extended Data Fig. 5.**
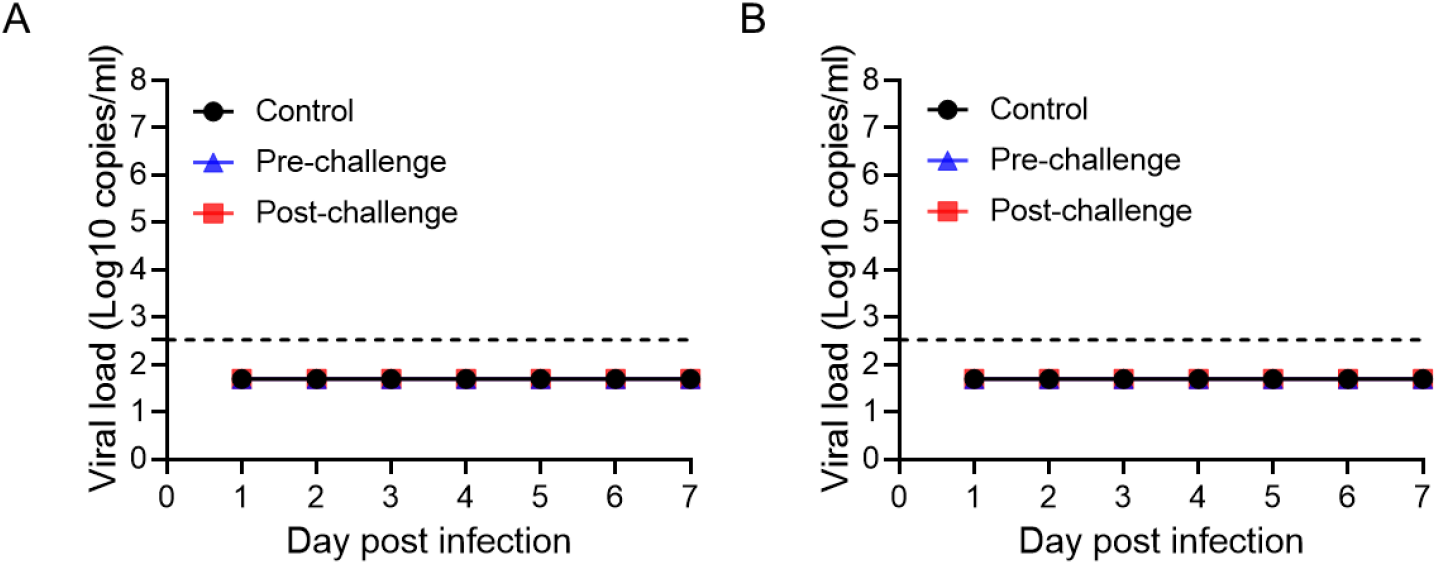
Viral titer of nasal swabs (A) or blood samples (B) of all monkeys were evaluated by qRT-PCR.

**Extended Data Fig. 6.**
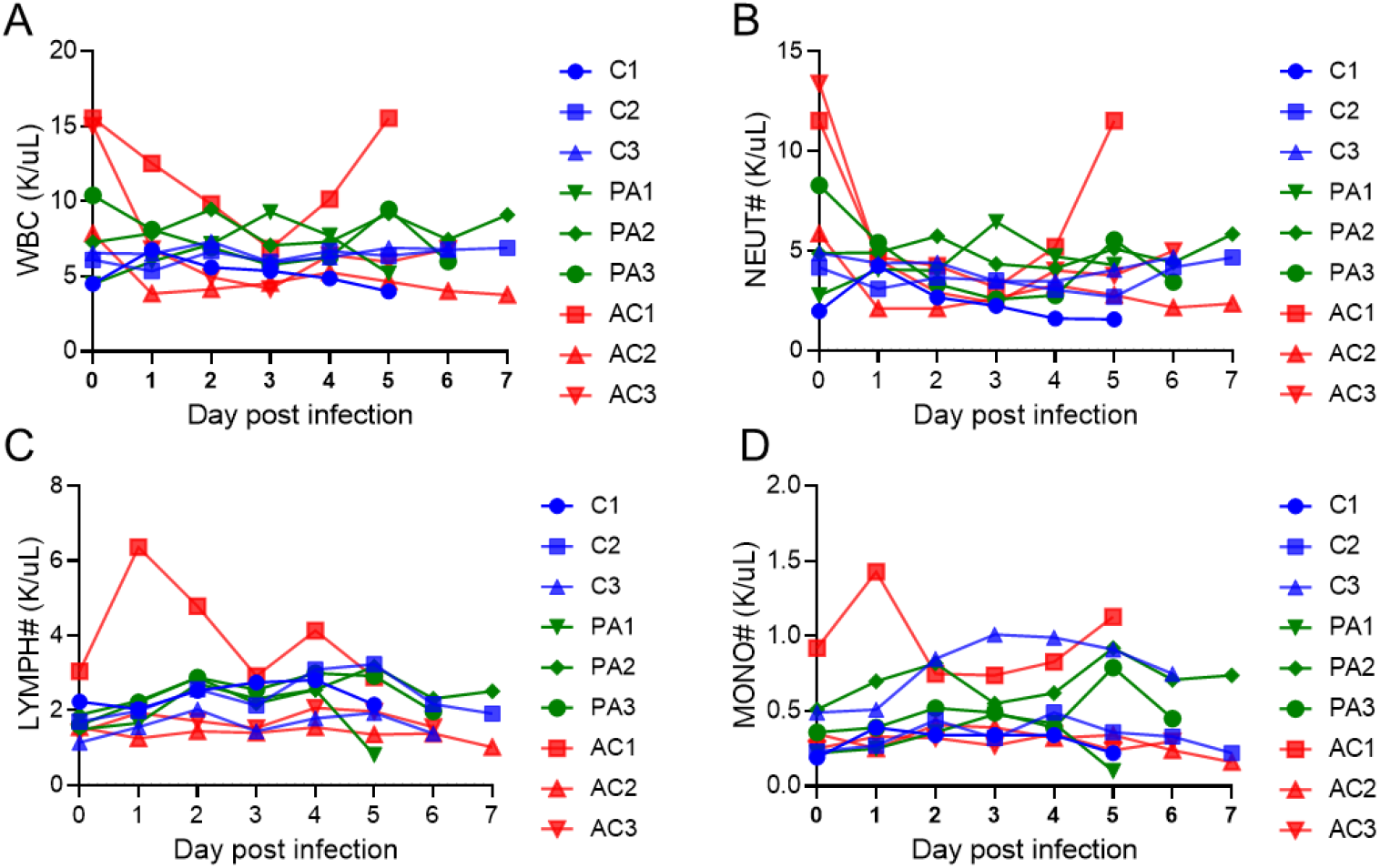
White blood cells (A), neutrophils (B), lymphocytes (C) and monocytes (D) in each monkey were monitored every day. “C” indicates the control group, “PA” indicates the pre-challenge group and “AC” indicates the post-challenge group.

